# Absolute quantification of cerebral metabolites using 2D ^1^H-MRSI with quantitative MRI-based water reference

**DOI:** 10.1101/2024.11.25.625224

**Authors:** Dennis C. Thomas, Seyma Alcicek, Andrei Manzhurtsev, Elke Hattingen, Katharina J. Wenger, Ulrich Pilatus

## Abstract

**Purpose:** Metabolite concentrations are valuable biomarkers in brain tumors (BT). However, correction of water relaxation effects often requires time-consuming quantitative MRI (qMRI) sequences on top of a lengthy spectroscopic water reference acquisition. The goal of this work was to develop and validate a fast metabolite quantification method where a 2D spectroscopic water reference acquisition is obtained using a fast qMRI protocol and single-voxel STEAM sequence.

**Methods:** A 2D sLASER sequence was acquired for MRSI. An 8-minute qMRI protocol was also acquired. A single-voxel unsuppressed water signal was acquired using a STEAM sequence. The H_2_O map, obtained from qMRI, was calibrated based on the STEAM-signal to obtain the spectroscopic water reference (proposed method). Five healthy volunteers and one BT patient were scanned at 3T. Concentrations obtained using the proposed and two reference methods, one where water relaxation effects were corrected using literature values (Reference method) and one where they were corrected using qMRI-derived values (Reference method with qMRI) were compared.

**Results:** In healthy subjects, WM metabolite concentrations obtained using water relaxation using literature values (Reference method) significantly differed from those using individual-specific corrections (Reference method with qMRI and proposed method). Bland-Altman analyses revealed a very low bias and SD of the differences between the ‘Reference method with qMRI’ and the ‘proposed-method’ (Bias<0.5% and SD<10%). The BT regions showed a ∼15% underestimation of metabolite concentrations using the ‘Reference method’.

**Conclusion:** For metabolite quantification, accurate water referencing with individual-specific corrections for water relaxation times was obtained in 8 minutes using the proposed method.

## Introduction

Proton Magnetic Resonance Spectroscopy (^1^H-MRS) is a powerful tool, capable of measuring *in vivo* metabolite concentrations^1^. In brain tumors (BT), changes in the relative and absolute concentrations of metabolites can serve as diagnostic and therapeutic biomarkers ^2–4^. In order to study the differences in the metabolite concentrations between patient and healthy cohort or between tumor tissue and healthy tissue in the same patient, the metabolite signal has to be referenced either to that of another metabolite or to that of water^5^. This is because, the signal amplitude at a certain region depends not only on the concentration of the metabolite, but also on factors which are hard to determine, like the receiver gain (B1-), the transmitter gain (B1+) and a scanner-dependent intensity scaling factor^5,6^. These factors could vary between regions, subjects and scanners. In the past, concentration ratios such as total Choline/total Creatinine (tCho/tCr) were widely reported for clinical use. However, this approach suffers from the ambiguity of attributing the changes to the metabolite of interest (in the numerator) or the reference metabolite (in the denominator) and the fact that small or inaccurate values for the denominator produce large variance in the metabolite ratios^7^. Another concern is the stability of the reference metabolite across age, gender, pathology or other factors^8^. Due to these reasons, it is advised that the metabolite concentrations be obtained by referencing the metabolite signal to that of water ^6,8–12^.

Semi-quantitative MRS can be achieved by calculating the ratio of each metabolite to the unsuppressed tissue water signal from the same voxel of interest (VOI), while assuming a standard MR-visible water content in the tissue^13^. However, for full quantification of metabolite concentrations with the internal water reference, it is necessary to adjust both water and metabolite signals for relaxation effects, account for tissue partial volume effects, and correct for regional variations of the water signal (in relation to B1 inhomogeneity and eddy currents) ^9^. Typically, the metabolite signal is acquired with a water-suppressed MRS sequence followed by the acquisition of the same MRS sequence without water suppression. The fitted metabolite signal intensity in a voxel is then divided by the fitted water signal intensity obtained from the same voxel, and relaxation effects of water and metabolites are corrected for with relaxation times obtained from literature^9^. However, this method of quantification has two disadvantages: First, the acquisition of both water suppressed and unsuppressed spectra under the same conditions nearly doubles acquisition time, which is undesirable for already lengthy 2D MRSI protocols. Second, using literature values of water relaxation may not be accurate, as the relaxation values in brain pathologies differ strikingly^14,15^. Even in healthy subjects, the water relaxation times show significant variations between subjects and brain regions of different tissue water density, making a single correction factor insufficient ^16,17^.

In order to address these inaccuracies, that are based on a number of assumptions, multiple studies have employed quantitative MRI (qMRI) to their quantification approach^18–22^. Notably, for 2D MRSI, Gasparovic et. al carried out a detailed quantification approach to correct for the water relaxation times by measuring the whole-brain H_2_O, T1 and T2 maps and correcting the acquired unsuppressed water signal for T1 and T2 relaxation^18^. The H_2_O map was used for obtaining molar concentrations of the metabolites from the molal concentrations (see equation 12 below). However, the total acquisition time just for obtaining an accurate relaxation-corrected water reference was close to 45 min, making this approach unfavorable for clinical applications.

The long acquisition time is partly due to the fact that both the unsuppressed water spectrum and the qMRI maps are acquired. However, as shown in the current work, this is a redundant approach since the spatial information encoded in the 2D MRSI unsuppressed water spectrum is also present in the qMRI maps, namely in the H_2_O and T1 maps^23–26^. This information obtained using qMRI techniques can be exploited to compute the water unsuppressed spectroscopic signal. Here, single-voxel unsuppressed STEAM acquisition was used to calibrate the H_2_O maps to the 1H MRSI data.

The aims of this work were:

1. To develop a fast, individual specific ^1^H-MRSI quantification protocol (∼8 min) without the need of an additional unsuppressed MRSI water reference, using qMRI and a single-voxel (SV) STEAM sequence for calibration.
2. To evaluate the differences between the obtained metabolite concentrations using the proposed method and currently recommended approaches (according to the consensus papers ^7,8,27^) in five healthy subjects and one BT patient.

### Theory

Consider that we are interested in measuring the concentration of a metabolite ‘met’ (*C*_*met*_). As outlined by Ernst et al.,^28^ *C*_*met*_ can be expressed as molal units (mol/kg of water) or molar units (mol/L of tissue). To elaborate, when expressed in molal units, *C*_*met*_ refers to the number of moles of ‘met’ present in 1 kilogram of ‘tissue water’, and when expressed in molar units, refers to the number of moles of ‘met’ present in 1 liter of ‘tissue’, which includes the non-water tissue and the Intracellular/Extracellular (IC/EC) tissue water. That is,

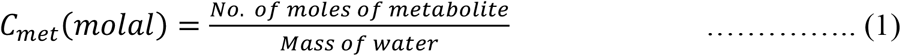

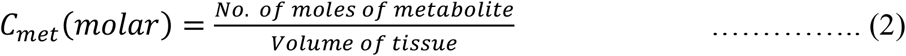

Note that, in this study, *C*_*met*_ excludes the CSF water, as most MRS detectable metabolites are not expected to be present in the CSF. The number of moles of a metabolite in each voxel is proportional to the metabolite signal, *S*_*met*_, in the spectra of the corresponding voxel of the acquired 2D MRSI data. In order to obtain absolute concentrations of metabolites, *S*_*met*_ needs to be calibrated, which, as mentioned in the ‘Introduction’ section, is usually achieved by recording the unsuppressed tissue water signal. As recommended in the 2020 consensus paper on preprocessing, analysis, and quantification in MRS,^8^ a water unsuppressed scan should ideally be recorded under the same conditions as the spectroscopic data. Relaxation corrections (T1, T2) for water can then be performed based on literature values (if segmented images are available) or based on the relaxation times measured using a qMRI protocol^18,19^. Since the approach with literature values is routinely used, it will be referred to as the “reference method” (Ref-method). The approach using values measured with qMRI will be referred to as the “reference method with qMRI” (Ref-method-with-qMRI). To avoid the lengthy acquisition of the spectroscopic water reference, we propose a third method which uses an unsuppressed SV STEAM sequence to calculate a spectroscopic water reference map from qMRI data. This method will be referred to as the “proposed method” (Proposed-method). Although most of the theory behind the Ref-method and Ref-method-with-qMRI have been explained in previous publications, we provide an overview of the theory for these approaches here, for sake of clarity.

#### Reference method (“Ref-method”)

In this approach, the unsuppressed water signal (S_*w*_) is obtained using the same 2D-MRSI sequence with the same parameters but without water suppression. It is therefore proportional to the number of moles of water in each voxel. By using the same sequence to acquire both the metabolites and water signals, the effect of the transmit and receive field inhomogeneities and other scaling factors are expected to be identical. However, there are two important corrections that need to be carried for absolute quantification: 1) T1 and T2 relaxation effects of water and metabolites and 2) correction for CSF partial-volume effects. The need for the first correction arises from the fact that in order to acquire data in reasonable acquisition times, a short TR (∼2s) is often used, and hence the signal is never fully relaxed. Further, hardware constraints and the sLASER sequence with 4 adiabatic full passage pulses limit the minimum achievable echo time, which leads to a non-zero T2 relaxation of the signal. The need for the second correction for CSF partial volume effects arises from the fact that the metabolites concentrations in CSF are usually negligible, hence, only the concentration of the metabolites in the non-CSF water are of interest. Including these corrections in the quantification gives us:

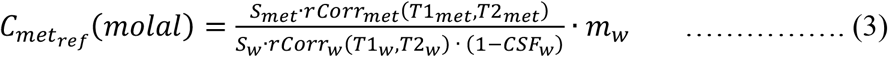

where, 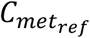 refers to the metabolite concentration obtained using the reference method. *rCorr*_*met*_ and *rCorr*_*w*_ refer to the relaxation correction factors for the metabolites and water relaxation respectively. T1_w_ and T2_w_ refer to the water T1 and T2 relaxation times, and T1_met_ and T2_met_ refer to the metabolite T1 and T2 relaxation times, which are obtained from literature for gray matter (GM) and white matter (WM) regions^29–33^. The molality of water (*m*_*w*_) is 55100 mmol/g. *CSF*_*w*_ refers to the CSF molal water fraction, as described by Gasparovic et. al^34^. In order to calculate *CSF*_*w*_, T1-weighted images are used to first compute WM and GM segmentations using SPM^35^. The segmentations provide f_WM_, f_GM_ and f_CSF_ which represent the volume fractions of WM, GM and CSF respectively. As described by Gasparovic et. al^34^. the molal water fraction of the CSF (*CSF*_*w*_) can then be obtained as follows:

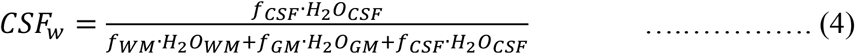

where, *H*_2_*O*_*WM*_, *H*_2_*O*_*GM*_ and *H*_2_*O*_*CSF*_ represent the relative molal water density in the WM, GM and CSF respectively, obtained from literature. Note that f_WM_, f_GM_ and f_CSF_ here represent the segmentations corrected for the difference in the point-spread function (PSF) between the high-resolution imaging data and the low-resolution MRSI data, as detailed in the Supporting Information. *rCorr*_*met*_ can be computed as follows:

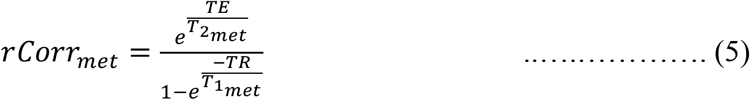

To compute *rCorr*_*W*_, the relaxation time correction factors for each of the different tissue compartments need to be computed separately, i.e,

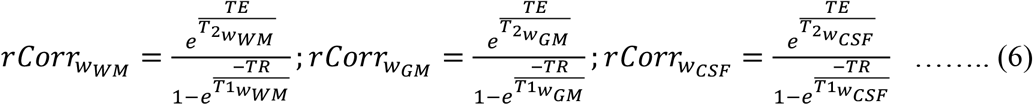

where, 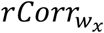 is the relaxation times correction factor for the tissue segmentation class *x. rCorr*_*w*_ can be then computed the sum of the product of the individual tissue segmentation class molal water fraction and the corresponding relaxation time correction factors:

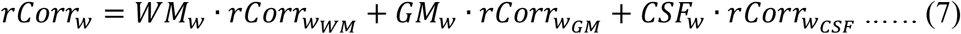

In order to compute 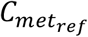 (*molar*), one can derive the equation similar to equation 5, but by using the tissue molar volume fractions instead of the tissue molal water fractions and replacing *m*_*w*_ with *M*_*w*_ (Molarity of pure water), as has been well described in Ref^36,37^. Here, however, we compute the metabolite molar concentration 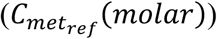 (excluding CSF) directly from 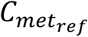(*molal*), by multiplying 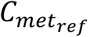(*molal*) by the water density in the voxel (excluding CSF):

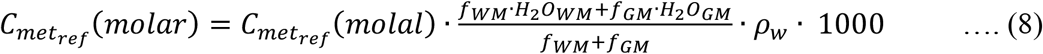

Here, *ρ*_*w*_ represents the density of pure water [g/L] and was assumed to be equal to unity. Equation 8 gives a direct relation between the molal concentrations (corrected for CSF partial volume) and the molar concentrations (corrected for CSF partial volume) when WM and GM segmentations are available.

#### Reference method with qMRI (“Ref-method-with-qMRI”)

In the ‘Ref-method’, parameters related to the water obtained from literature are: 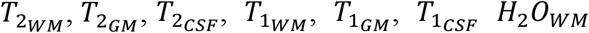 and *H*_2_*O*_*GM*_. In the ‘Ref-method-with-qMRI’, these parameters are measured using qMRI sequences for T2 mapping, T1 mapping, and H_2_O mapping relying on vendor-based sequences^29^. The parameters of acquisition for the qMRI sequences are described in the methods section. In equation 4, for the Ref-method-with-qMRI, the computation of *CSF*_*w*_ and *rCorr*_*w*_(*T*1, *T*2) changes. In this case, the molal water fraction of the CSF obtained using ‘Ref-method-with-qMRI’ 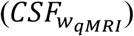 is computed using the water content map (*H*_2_*O*_*map*_):

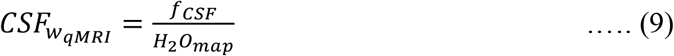

*rCorr*_*w*_ obtained using ‘Ref-method-with-qMRI’ 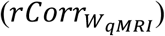, is computed using the T1 and T2 maps (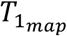 and 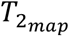), as follows:

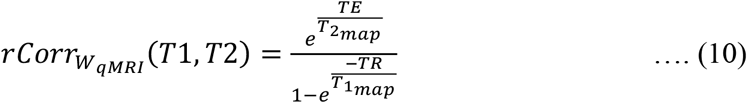

The final metabolite concentrations are then calculated according to the following equations (similar to equations 5 and 10):

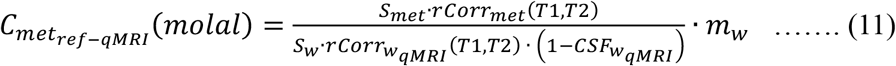

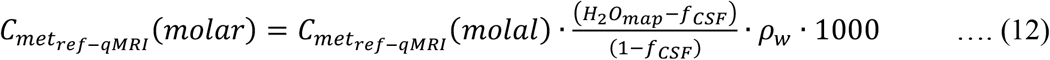

Note that the 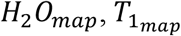 and 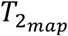 are corrected for the difference in PSF due to the differing resolution between MRSI and imaging as detailed in the Supporting Information.

#### Proposed method (“Proposed-method”)

The proposed method for absolute quantification of metabolites is graphically depicted in **Figure 1**. T1, T2*, B1+ and H_2_O maps (*T*1_*map*_, *T*2^∗^_*map*_, *B*1_*map*_ & *H*_2_*O*_*map*_) are obtained using the same vendor-based qMRI protocol used for the ‘Ref-method-with-qMRI’^29^. For calibration of the spectroscopic data, a water-unsuppressed SV STEAM sequence is run with the voxel positioned in the normal appearing WM (NAWM) (voxel size = 10^3^ mm^3^). This measured water signal is labeled 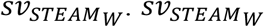 is corrected for the B1+ inhomogeneity, T1 relaxation and T2 relaxation using the B1+, T1 and T2* values of that voxel obtained from the mp-Qmri maps. Since the fast qMRI protocol used does not produce a *T*2_*map*_, the T2 value of the water in the STEAM voxel is estimated using the T2* map: 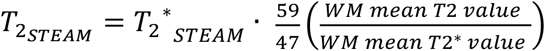. Further, 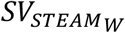. is corrected for the difference in the voxel size between STEAM and sLASER.

**Figure 1:**
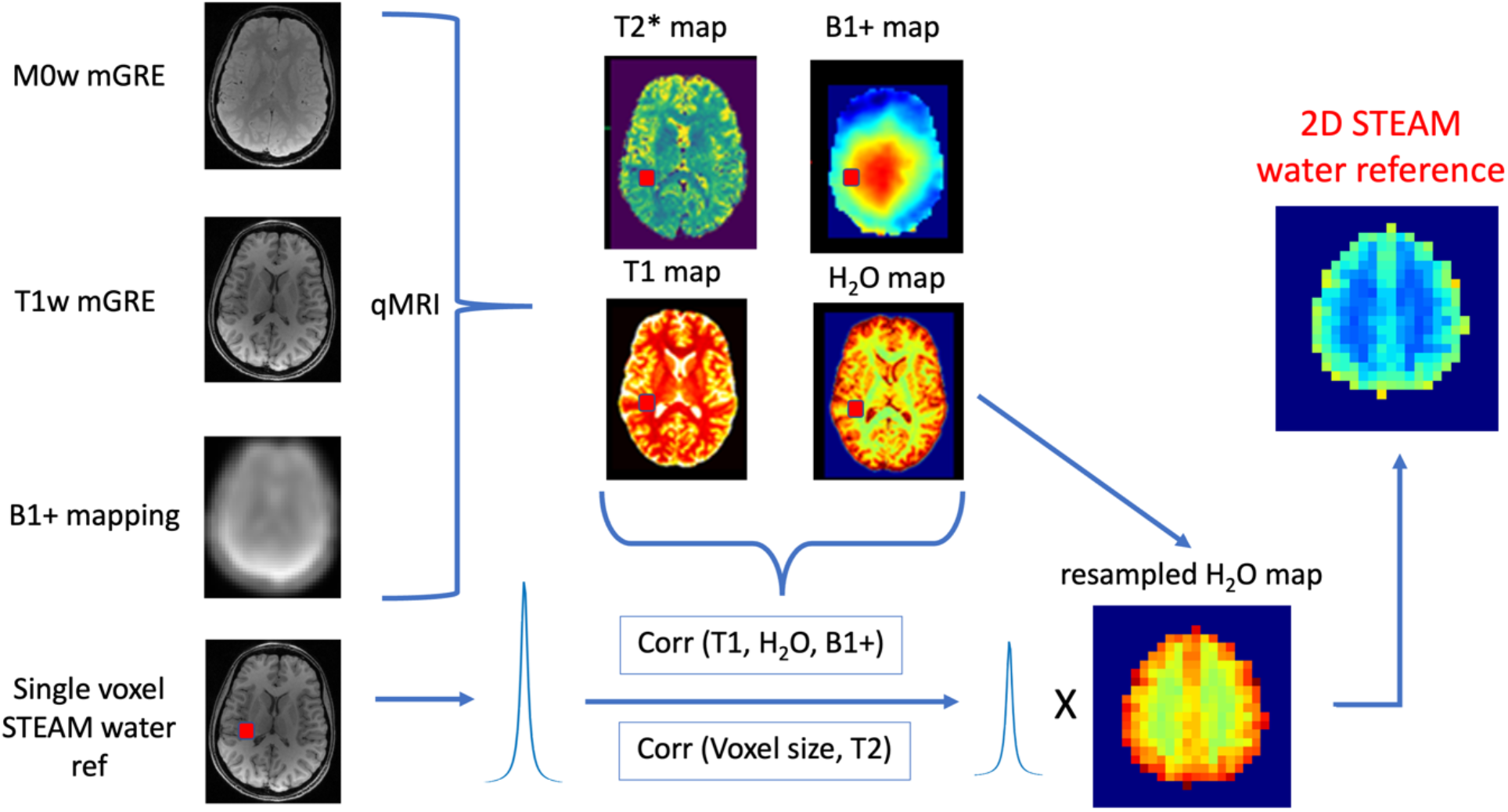
Graphical depiction of the Proposed-Method. For the qMRI protocol, two multi-echo gradient-echo (mGRE) images (M0w/T1w) images and two EPI images (45°/90°) are obtained. A single-voxel water unsuppressed STEAM signal is obtained from a voxel placed in the NAWM. The STEAM water signal is then corrected for T1, T2, B1+ and voxel size, to obtain the corrected water signal which is then used to calibrate the *H*_2_*O*_*map*_ which yields the 2D STEAM water reference 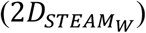.

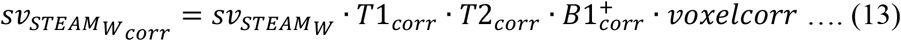

where,

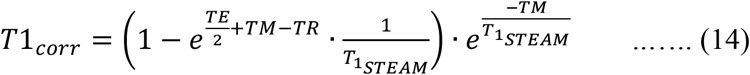

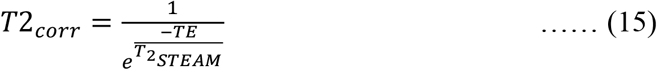

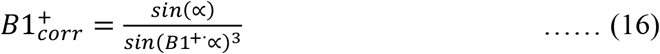

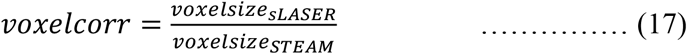

Here, 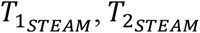 and 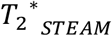 refer to the T1, T2 and T2* of water in the STEAM voxel. ∝ refers to the nominal flip angle, TM refers to the mixing time.

Finally, 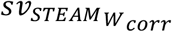 is used to calibrate the *H*_2_*O*_*map*_, extending the SV measurement to obtain a calibrated 2D STEAM water reference 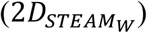:

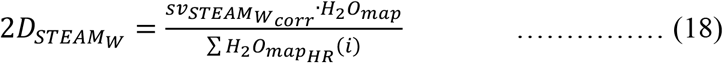

where *i* refers to the voxels in the high-resolution 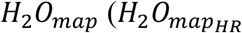 which lie within the 10^3^ mm^3^ STEAM voxel. 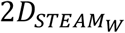 can then be used as the spectroscopic water reference (*S*_*w*_) and the absolute concentration of the metabolites can be computed in the same way as described in the ‘reference method with qMRI’ method, i.e equations 9-12.

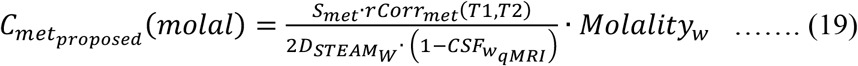

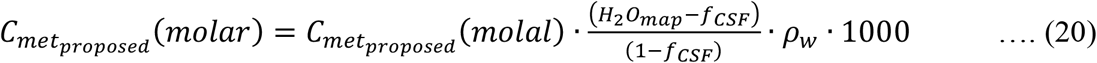

To summarize, the two differences between the Ref-method-with-qMRI and the Proposed-method are:

1. The T2 map for the Proposed-method is not directly acquired. The T2 value of the STEAM voxel is estimated from the T2* value in the STEAM voxel using a constant correction factor (1.255).
2. The spectroscopic unsuppressed water reference is not directly acquired for the proposed method, but instead calculated by acquiring a SV STEAM sequence and calibrating the H_2_O map using the STEAM signal.

## Methods

### Study protocol

#### MRS

*In vivo* measurements were performed on five healthy subjects and one BT patient on a whole-body 3T MR Scanner (MAGNETOM Prisma (VE11C), Siemens Healthineers, Erlangen, Germany). The MRI sequence protocol was the same for the healthy subjects and the BT patient. MRSI was acquired with a 2D sLASER sequence^38^ with acquisition parameters as detailed in **Table 1**. For the ‘Ref-method’ and ‘Ref-method-with-qMRI’, the spectroscopic water reference was acquired with the same parameters as the 2D sLASER, but without water suppression. For the Proposed-method, a 10 mm^3^ SV STEAM sequence was acquired in the NAWM, with no water suppression.

**Table 1:** Acquisition parameters of the spectroscopic sequences.

|  | 2D sLASER | 2D sLASER water reference | SV STEAM |
| --- | --- | --- | --- |
| TR | 2000 ms | 2000 ms | 10000 ms |
| TE | 40 ms | 40 ms | 20 ms |
| TM | - | - | 10 ms |
| FA | 90 | 90 | 90 |
| BW | 2000 Hz | 2000Hz | 1200Hz |
| Interpolated Resolution | 7.5 x 7.5 x 12 mm<sup>3</sup> | 7.5 x 7.5 x 12 mm<sup>3</sup> | 10 x 10 x 10 mm<sup>3</sup> |
| Matrix size | 20 x 20 | 20 x 20 | - |
| Averages | 2 | 1 | 1 |
| In-plane FOV | 240 x 240 mm<sup>2</sup> | 240 x 240 mm<sup>2</sup> | - |
| No. of slices | 1 | 1 | - |
| Slice thickness | 12 | 12 | - |
| Water suppression | CHESS | - | - |
| TA | 10:52 min | 8 min | 10s |

For the five healthy subjects, the spectroscopic slice was positioned above the corpus callosum, so that no part of the lateral ventricles was within the Volume of Interest (VOI). For the brain tumor patient, the VOI was positioned to cover the solid part of the tumor and the contralateral healthy tissue.

#### qMRI

For the Ref-method-with-qMRI and Proposed-method, an 8-minute vendor-based protocol for obtaining whole-brain T1, H_2_O, T2* and QSM maps was acquired with the protocol described <u>by</u> Thomas et. al.^29^. This included measuring 2 multi-echo Gradient-Echo (mGRE) images (M0w and T1w), and two EPI images for B1 mapping (**Figure 1**). Additionally, for the Ref-method-with-qMRI, T2 mapping with three single-echo spin-echo images and 20 slices (to reduce acquisition time) was acquired. The slices were positioned to cover the VOI of the MRSI slab. Acquisition parameters are detailed in **Table 2**.

**Table 2:**
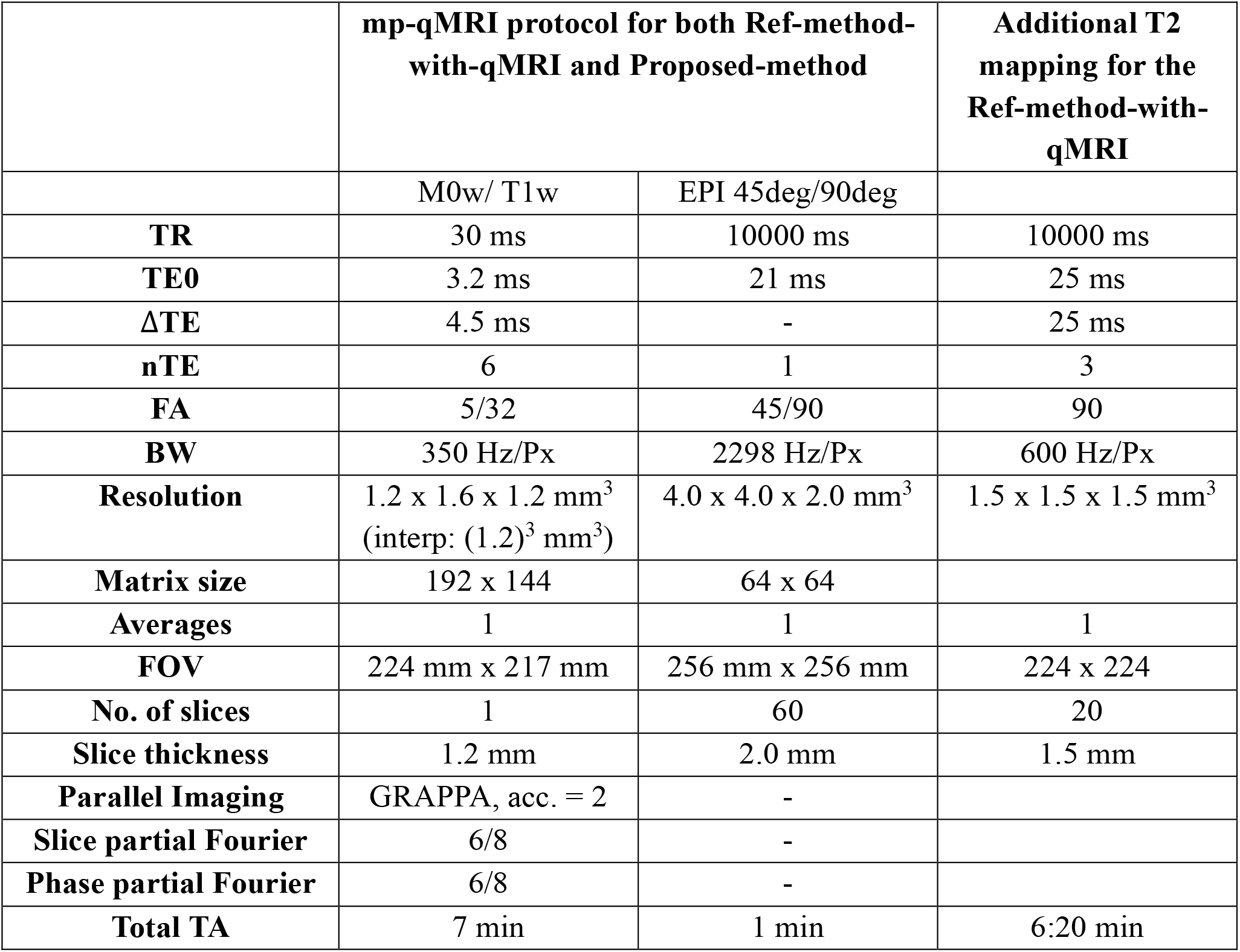
Acquisition parameters of qMRI sequences.

|  | **mp-qMRI protocol for both Ref-method-with-qMRI and Proposed-method** |  | **Additional T2 mapping for the Ref-method-with-qMRI** |
| --- | --- | --- | --- |
|  | M0w/ T1w | EPI 45deg/90deg |  |
| **TR** | 30 ms | 10000 ms | 10000 ms |
| **TE0** | 3.2 ms | 21 ms | 25 ms |
| **ΔTE** | 4.5 ms | - | 25 ms |
| **nTE** | 6 | 1 | 3 |
| **FA** | 5/32 | 45/90 | 90 |
| **BW** | 350 Hz/Px | 2298 Hz/Px | 600 Hz/Px |
| **Resolution** | 1.2 x 1.6 x 1.2 mm<sup>3</sup><br>(interp: (1.2)<sup>3</sup> mm<sup>3</sup>) | 4.0 x 4.0 x 2.0 mm<sup>3</sup> | 1.5 x 1.5 x 1.5 mm<sup>3</sup> |
| **Matrix size** | 192 x 144 | 64 x 64 |  |
| **Averages** | 1 | 1 | 1 |
| **FOV** | 224 mm x 217 mm | 256 mm x 256 mm | 224 x 224 |
| **No. of slices** | 1 | 60 | 20 |
| **Slice thickness** | 1.2 mm | 2.0 mm | 1.5 mm |
| **Parallel Imaging** | GRAPPA, acc. = 2 | - |  |
| **Slice partial Fourier** | 6/8 | - |  |
| **Phase partial Fourier** | 6/8 | - |  |
| **Total TA** | 7 min | 1 min | 6:20 min |

### Data processing

#### Metabolite spectra fitting

Spectroscopic data were processed using the LCModel software^39^. Metabolite basis sets were simulated for the sLASER pulse sequence at TE = 40 ms using the jMRUI plug-in NMR-ScopeB (version 6.0, available at http://www.jmrui.eu^40^) with prior knowledge of chemical shifts and J-coupling. In the control file used for LCModel processing, the parameters ‘atth2o’, ‘attmet’ and ‘wconc’, responsible respectively for correction of the water relaxation effects, metabolite relaxation effects, and water density, were set equal to 1, avoiding correction of water and metabolite relaxation effects and water density. The re-slicing and averaging of anatomical and quantitative data into the spectroscopic domain and all the quantification steps were carried out in Python v3.9.12, using home-written scripts.

#### Quantification of metabolite concentrations

T1, H_2_O, T2* and QSM mapping were carried out with the post-processing steps detailed by Thomas et. al^29^. For T2 mapping, the three SE images acquired at echo times of 25 ms, 50 ms and 75 ms were voxel-wise fitted using the following equation:

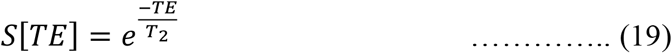

where, *S*[*TE*] denotes the signal at echotime TE.

For the Ref-method the unsuppressed sLASER water reference scan was given as the water reference. LCModel computes the concentration of the metabolites by normalizing the metabolite signal to this water signal. The LCModel output corresponds to 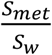. Then, the corrections for water relaxation, metabolite relaxation and CSF partial volume effect were carried out as detailed in the Theory section. The literature values used for the water and metabolite relaxation correction are given in **Supporting Information Table 1**. The first-echo image of the T1-weighted mGRE was used for the WM, GM and CSF segmentation using SPM. For the Ref-method-with-qMRI, the corrections were carried out as detailed in equations 11 and 12. Instead of using values from literature for T1, T2 and water content (H_2_O) of WM, GM, and CSF, the actual measured T1, T2, and H_2_O maps were used. CSF segmentation was performed for correction of CSF partial volume effect as carried out in equation 11. For the Proposed-Method, the 2D spectroscopic water reference was obtained using qMRI and STEAM acquisition. The STEAM unsuppressed water signal was propagated to form a 2D STEAM array of the same matrix size as that of the sLASER metabolite acquisition and given as the water reference in LCModel. The metabolite concentrations that were obtained from LCModel were then corrected using equations 13-20.

### Statistics

Since metabolite concentrations show regional variations in the brain, four VOIs were defined in order to compare the methods in the healthy volunteers: ‘Anterior’, ‘Posterior’, ‘Left’ and ‘Right’. The VOIs were manually defined on the MNI-152-brain T1-weighted image in the MNI space^41^. The VOIs were then transformed into the subject native space for each of the five volunteers using FSL, and analyses were carried out. In each of these VOIs, the concentrations of total N-Acetyl Aspartate (tNAA), tCr and tCho, obtained using the three methods were compared. Non-parametric Friedman’s repeated-measures ANOVA (rm-ANOVA) was used to assess the significance of the difference between the methods, with post-hoc Durbin-Conover tests carried out to determine the groups with significant differences. This was carried out for each of the VOIs and each of the metabolites separately. Bland-Altman analysis was performed for the Ref-method-with-qMRI and the Proposed-method, as well as for the Ref-method-with-qMRI and the Ref-method.

## Results

**Figure 2** shows the metabolite maps obtained using the three methods. Upon visual inspection of the metabolite maps, the Ref-method-with-qMRI and Proposed-method maps showed good agreement. The Ref-method maps on the other hand differed noticeably, with higher metabolite concentrations in the non-WM regions, in comparison to the Ref-method-with-qMRI and Proposed-method maps.

**Figure 2:**
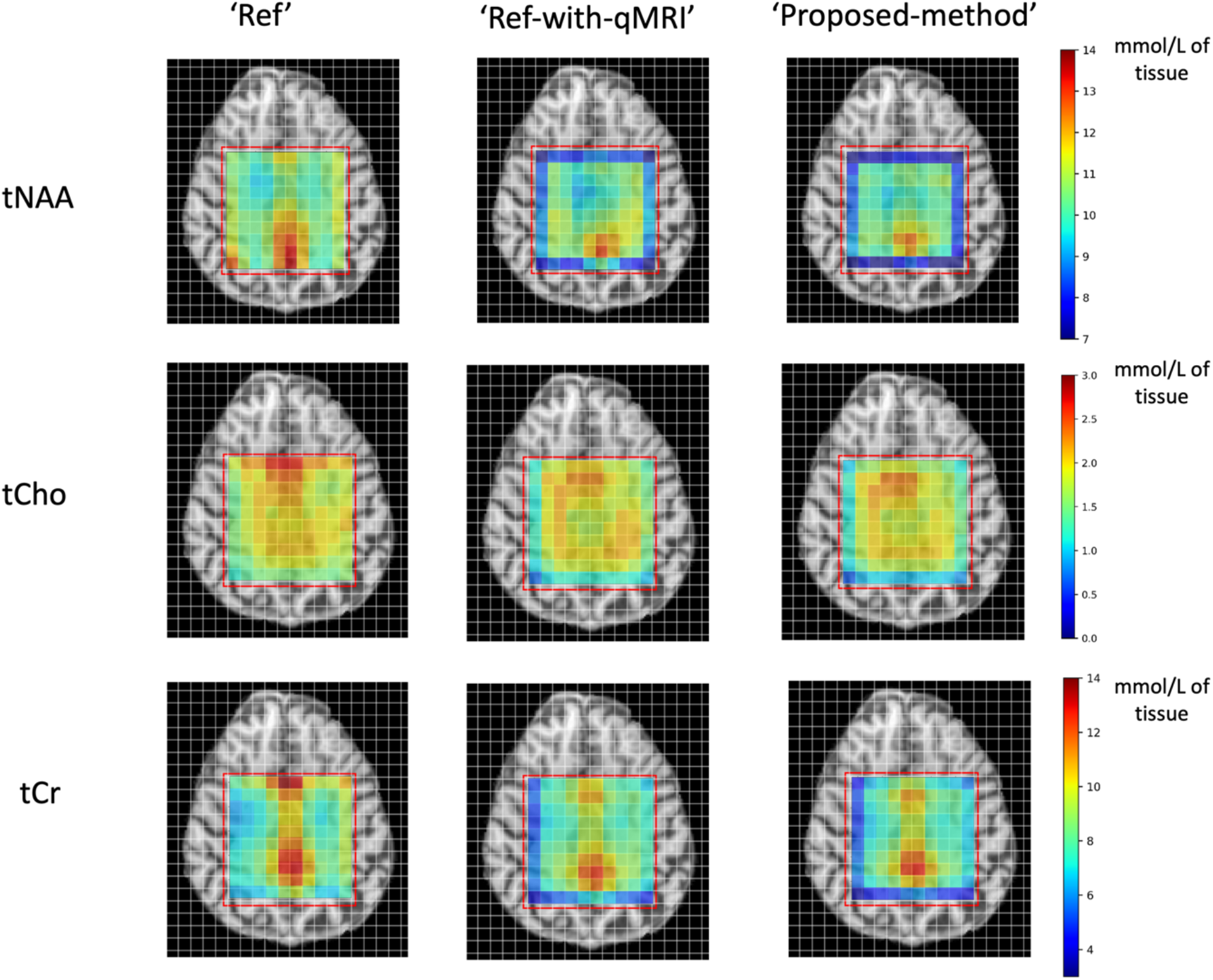
Molar concentration metabolite maps of Subject 2 obtained with the three methods. The Ref-method-with-qMRI and Proposed-method agree well with each other. The Ref-method maps qualitatively differ slightly from the Ref-method-with-qMRI maps. The red box denotes the spectroscopic ROI.

**Figure 3** shows bar plots for mean ‘**tNAA**’, ‘**tCr**’ and ‘**tCho**’ values across the five healthy subjects, as well as the four VOIs, defined in the MNI space (upper left). In the Anterior and Posterior VOIs, where GM and CSF are predominant, the metabolite values did not differ between the three method (p-values displayed in **Table 3**). In the Left and Right ROIs however, which predominantly consist of WM, the Ref-method values were significantly lower than the values obtained using the Ref-method-with-qMRI (**Table 3**). There were no significant differences, however, between metabolite concentrations obtained with the Proposed-method and those obtained with the Ref-method-with-qMRI. Metabolite concentrations derived from each of the three methods and the defined ROIs are presented in **Table 4**.

**Table 3:** p-values for pooled metabolite concentration differences between the Ref-method, Ref-method-with-qMRI and proposed method across the five healthy subjects.

|  | tNAA | tCho | tCr |
| --- | --- | --- | --- |
| ROI-Anterior | p = 0.549 | p = 0.449 | p = 0.449 |
| ROI-Posterior | p = 0.449 | p = 0.247 | p = 0.449 |
| ROI-Left | p < 0.01 | p < 0.01 | p < 0.01 |
| ROI-Right | p < 0.01 | p < 0.01 | p < 0.01 |

**Table 4:**
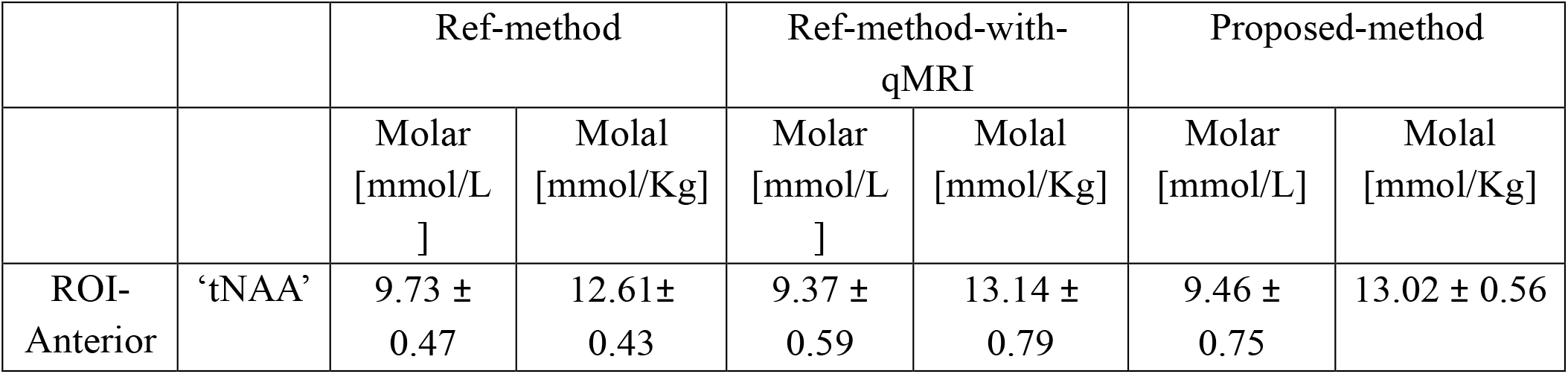
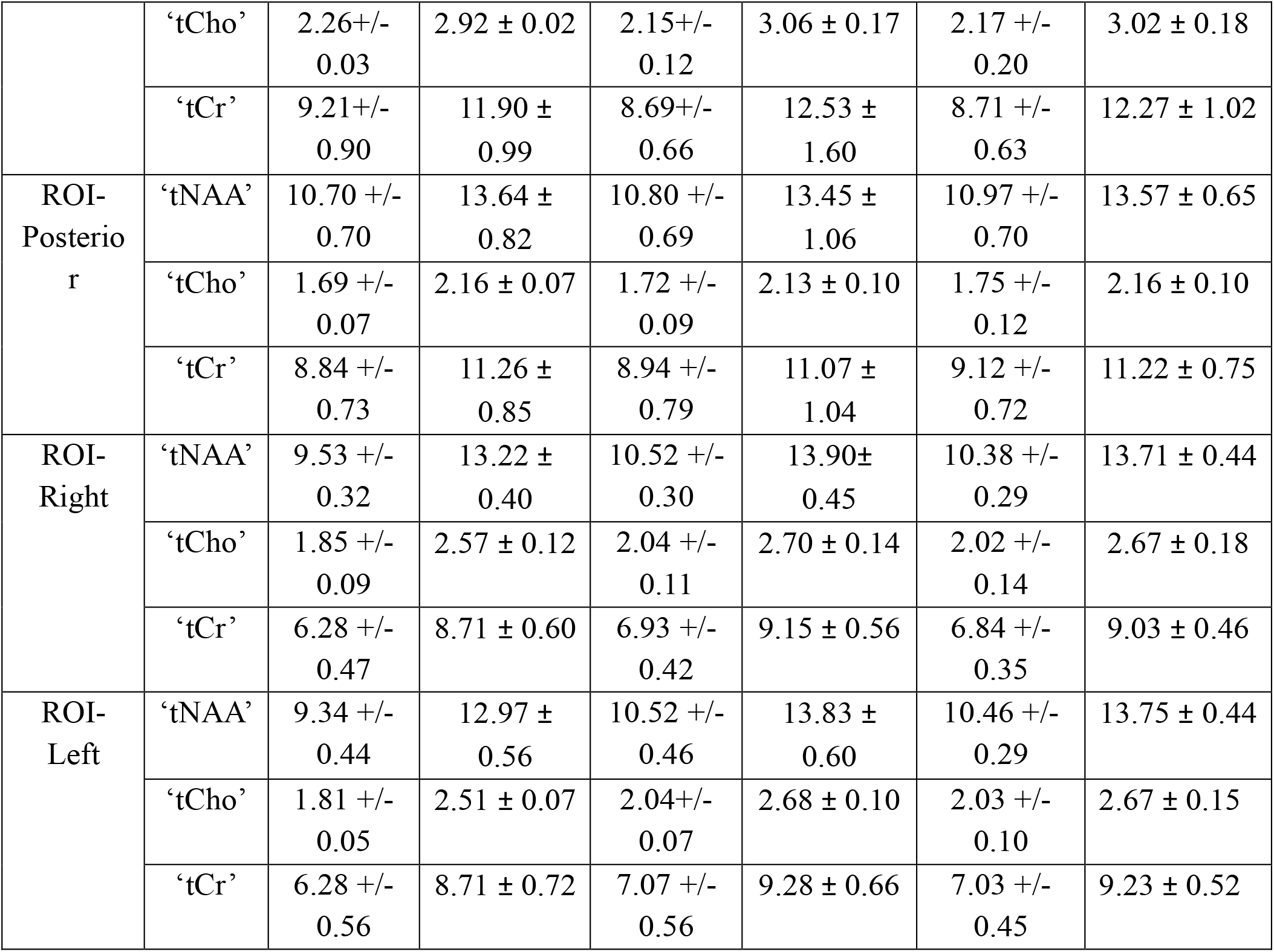
Mean Molar and Molal metabolite concentrations in ROIs obtained using the three methods.

**Figure 3:**
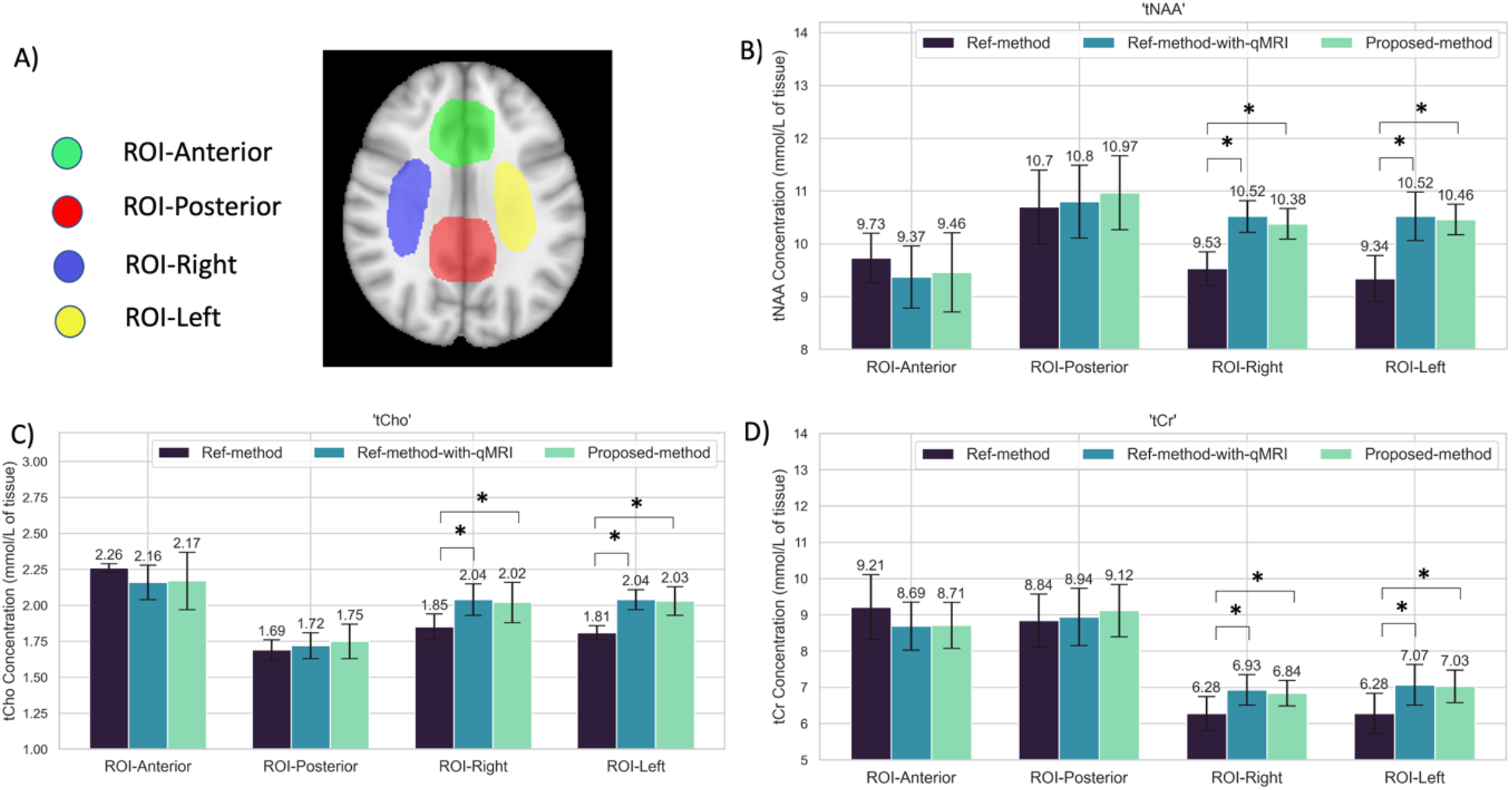
A) The ROIs defined on the MNI template. B, C, D) Box plots showing the mean metabolite molar concentrations averaged across the five subjects for the different ROIs for tNAA, tCho and tCr, respectively. The error bars denote the standard deviation across the five healthy subjects.

Voxels located on the outer borders of the ROI in 2D-MRSI are commonly deemed unreliable because of the imperfect slice profile and ignored in the analysis. Interestingly, as noticeable in **Figure 2**, the bordering voxels reveal differences between the three methods, with a notable underestimation of the metabolite values obtained using the Ref-method-with-qMRI and the Proposed-method, and a slight overestimation of metabolite values in the Ref-method.

**Figure 4** shows the results of the Bland-Altman analysis for the comparison of the Ref-method-with-qMRI and Proposed-method, as well as the Ref-method-with-qMRI and Ref-method. Regarding interchangeability of the methods, the Ref-method-with-qMRI and Proposed-method were in agreement with a small bias of <0.5% and a standard deviation (SD) of <10% of the mean metabolite concentrations. The differences between Ref-method-with-qMRI and Ref-method showed a larger Bias (∼5%) and SD (∼20%) in comparison to the differences between Ref-method-with-qMRI and Proposed-method.

**Figure 4:**
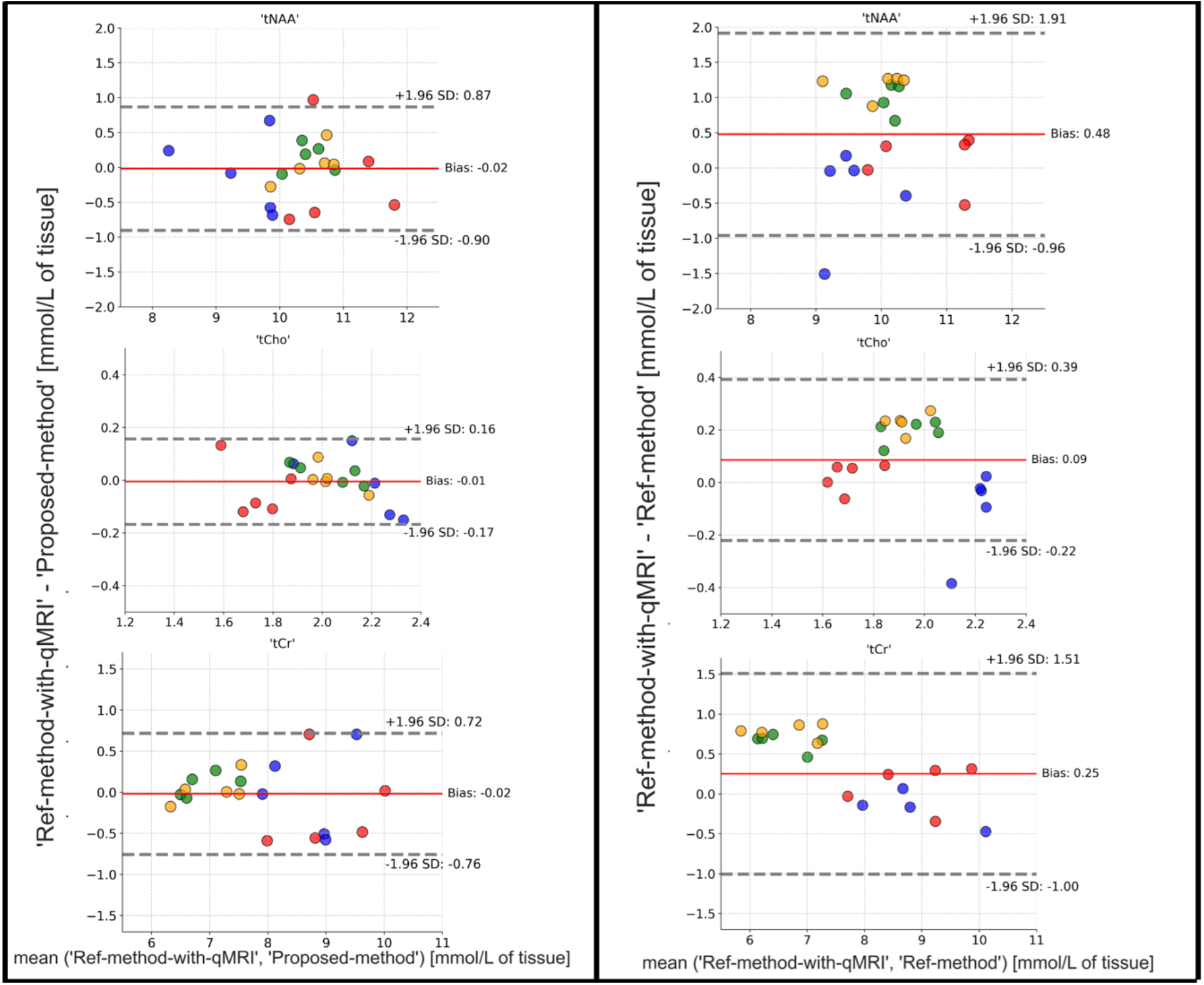
Bland-Altman analyses. Left Column: Difference between the Ref-method-with-qMRI and the Proposed-method are plotted on the y-axis; x-axis shows the mean of the metabolite concentrations obtained using the two methods. Right Column: Difference between the Ref-method-with-qMRI and Ref-method are plotted on the y-axis; x-axis shows the mean of the metabolite concentrations obtained using the two methods. Data points by color: Green = ROI Anterior, Red = ROI Posterior, Blue = ROI Right, Yellow = ROI Left

**Figure 5** shows the results of application of the three quantification methods in a BT patient. Qualitatively, the maps reveal differences between the Ref-method and the Proposed-method. Difference maps were calculated for quantitative display. The difference maps reveal an overestimation of the metabolite concentrations in the healthy tissue and underestimation of the metabolite concentrations in the tumor tissue with the Ref-method in comparison to the Proposed-method. In the tumor region, the Ref-method metabolite concentrations are ∼15% lower than those obtained using the Proposed-method.

**Figure 5:**
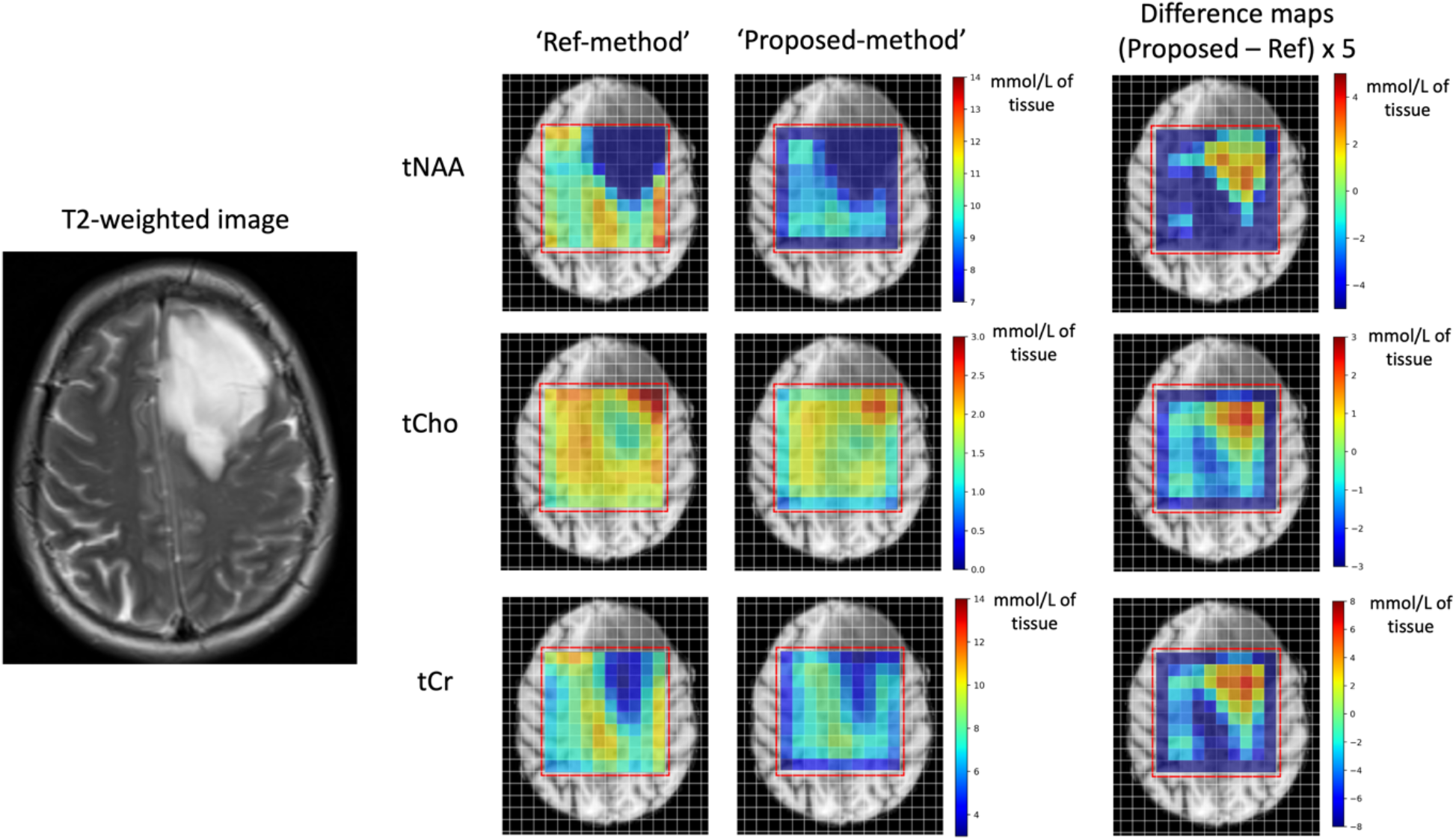
Metabolite concentration maps in a BT patient (Diagnosis: astrocytoma, IDH-mutant, WHO grade 3). The T2-weighted image shows an intra-axial BT in the left frontal lobe. The metabolite maps obtained using the Ref-method and the Proposed-method are shown here. As expected, both methods show a decrease in tNAA and tCr, and an increase in tCho concentrations in the tumor area. The difference maps between the two methods (multiplied by 5) are displayed on the right. In the tumor regions, there is an underestimation of the metabolite values obtained using the Ref-method. For improved visualization, values of the difference map are multiplied with 5 (5 mmol/L∼1 mmol/L).

## Discussion

In this work, we propose a fast, accurate, and easily implementable qMRI-based approach for absolute quantification of cerebral metabolites measured with a 2D MRSI sequence. The required mGRE, EPI and SV STEAM sequences are sequences offered by all vendors.

In comparison to other efforts employing qMRI for individual-specific metabolite quantification, our approach is faster for two reasons:

1. a fast protocol for qMRI is used to correct for water relaxation times
2. the measurement of the 2D unsuppressed water reference (∼8 min) is avoided, replacing it with the H2O map obtained from the qMRI protocol and calibrating it using a fast SV STEAM sequence (10 s).

Measuring qMRI enables a more accurate correction of water relaxation, obtaining the true water content of the tissue for calculation of metabolite concentration, and for replacement of a long redundant measurement of 2D unsuppressed water reference (∼8min) with a fast single voxel STEAM sequence (10s). The ‘Proposed-method’ does not rely on T2 mapping, as only the T2 value in the STEAM voxel (preferably placed in NAWM) needs to be known. The T2 value of the STEAM voxel is estimated from the T2* value obtained with the qMRI protocol. We hypothesized that this estimation is sufficient for the purpose of metabolites quantification. Our hypothesis was confirmed by the excellent agreement between the ‘Proposed-method’ and the ‘Ref-method-with-qMRI’ that is based on an additional full T2 mapping.

Since the sLASER sequence uses adiabatic pulses, the metabolite signal was considered to be insensitive to transmit profile (B1+) inhomogeneities^38^. For the STEAM measurement, the B1+ value was obtained from qMRI. If sequences that do not rely on non-adiabatic pulses (e.g., a PRESS sequence) are used, a correction for B1+ inhomogeneities for the metabolite signal should be carried out. This correction is readily available from the qMRI protocol used. sLASER was chosen in this study, as it has been shown that sLASER leads to a significant reduction of Chemical Shift Displacement artefacts in comparison to PRESS and STEAM. With respect to the receive field inhomogeneities, an in-line correction option provided by the vendor (“Prescan normalize”) was used. Thus, the spectroscopic data obtained from the scanner were already corrected for receive field inhomogeneities. If a vendor-based normalization option is not available, this correction would need to be performed offline. Ideally, the bias field maps generated as part of the post-processing step in water content mapping could be used, as described by Lecocq et. al ^19^.

Using the ‘Ref-method’ led to an underestimation of metabolite concentrations in ROIs with predominantly WM (ROI-Left and ROI-left) in the five healthy subjects, in comparison to the ‘Ref-method-with-qMRI’. ROIs containing predominantly of GM and CSF (ROI-Anterior and ROI-Posterior) showed no significant difference in the metabolite concentrations obtained using the three methods. This might be a result of the increased uncertainty of the literature and measured qMRI values in CSF voxels or voxels affected by CSF partial volume effects. Even in GM, literature T2 values range from 55ms to 110ms, making the metabolite concentrations obtained using the Ref-method highly sensitive to the choice of the GM T2 value. The GM T1 and water content values also have a higher SD across subjects and a broader within-subject distribution within in comparison to WM. These uncertainties in the GM/CSF-predominant ROIs lead to an increased SD and a decreased statistical power to detect biases. In WM on the other hand, qMRI is relatively well-behaved, and shows significantly lower standard deviations. The literature values of T1, T2 and H_2_O also fall in a narrower range. The lower error of qMRI mapping and correction can lead to an increased statistical power in the ROIs with predominantly WM. This is also evident in the lower inter-subject SD of the metabolite values in ROI-Left and ROI-Right as compared to ROI-Anterior and ROI-Posterior (**Figure 3** and **Table 3**). VFA-based T1 mapping and T2 mapping are known to be less accurate in CSF. In voxels affected by CSF partial volume, the ‘Proposed-method’ has the advantage of not using the T1 maps directly, but instead the final water content map, which is corrected for T1 and T2* relaxation in the WM and GM. A constant T1 value of 4300ms is used for correction of the T1 relaxation of the CSF voxels and further, in the water content maps, CSF voxels are normalized to a water content value of 1. In summary, the Proposed-method is less sensitive to inaccuracies of CSF T1/T2 mapping.

The discrepancy between the ‘Ref-method’ and ‘Ref-method-with-qMRI’ metabolite values highlights the bias resulting from using literature T1 and T2 values for quantification and further underlines the importance of actual relaxation time measurements, even in healthy subjects. In healthy subjects, regional variations in the T1, T2 and H_2_O are well known^42^. Using global T1, T2 and H_2_O literature values leads to a bias that is confirmed in our study. Even at the relatively short echo-time of 40 ms used in this study, differences in the metabolite concentrations are observed when relaxation correction was carried out using qMRI. At longer echo-times these differences are only expected to increase, and the correction for water relaxation time using measured relaxation parameters will be all the more crucial in order to obtain accurate metabolite concentrations. In BT patients, segmentation maps usually classify the tumor regions as GM voxels due to increased water content in tumor regions. Using normal GM literature values for water relaxation correction in BT regions, naturally leads to an underestimation of the local metabolites, which is depicted in the difference maps in **Figure 5**. Since BT can be highly heterogeneous, it would be impossible to accurately correct for water relaxation effects without using qMRI sequences. In the IDH-mutant astroctyoma patient scanned in this study, the BT regions revealed a ∼15% difference between the metabolite values obtained using the ‘Ref-method’ and the ‘Proposed-method’.

**Figure 6** shows the advantage of the method proposed in this study. Apart from the metabolite maps (in both molal and molar concentrations), four high resolution qMRI maps are obtained with inherent value in BT patients ^43–45^. In BT, measuring the actual relaxation times is a necessity for absolute metabolite quantification due to the heterogenous increase in T1, T2, and water content values^46^, depending on the tumor type and morphology.

**Figure 6:**
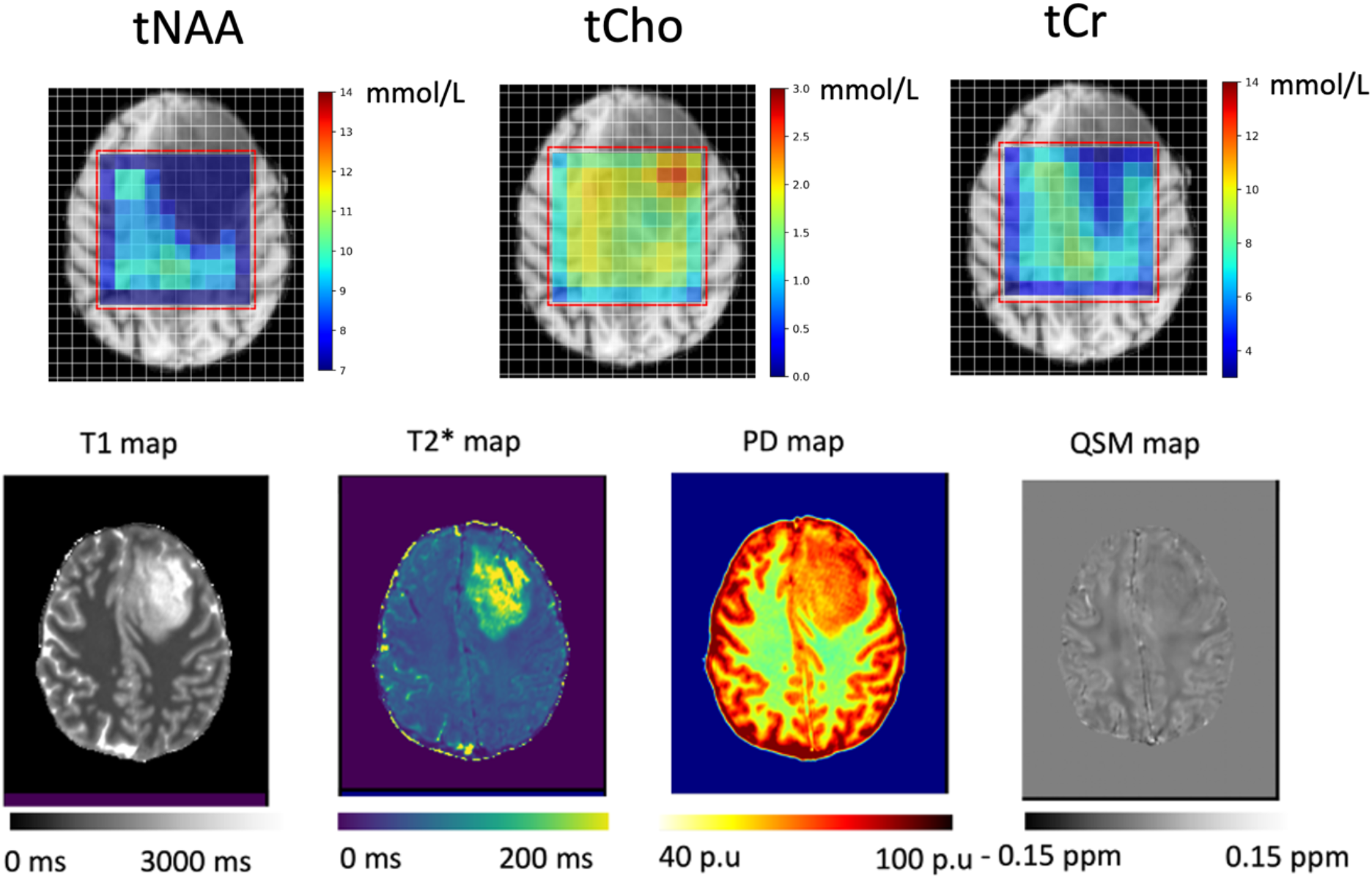
Final outputs of the absolute quantification approach using the Proposed-method are displayed here. Upper row: Metabolite concentration maps with individual-specific correction for water relaxation effects (‘tNAA’, ‘tCho’ and ‘tCr’ metabolite maps are shown). Bottom row: The four qMRI maps (T1, T2*, H_2_O and QSM) obtained with the mp-qMRI protocol are displayed.

## Limitations

One caveat in the proposed method is the underestimation of the water content in large BT ^47^. It has been shown that water content mapping techniques underestimate the water content in large BT, because the bias field correction algorithms mistakenly attribute the increased intensity in BT regions to that arising due to receive field inhomogeneities. In the current study, this effect is not accounted for. Future studies are needed to develop novel strategies for accurate correction of the receive field inhomogeneities in large BT. A second limitation of the study is that the metabolite relaxation was corrected for using the relaxation times derived from literature. Ideally a complete absolute quantification protocol should also include metabolite relaxation time measurements too. However, this is a more complex problem leading to very long acquisition times for the quantification protocol. Fast mapping of metabolite relaxation times is an area of active research and further work is needed in this direction. A third limitation of the proposed method is that although it enables an accurate quantification of metabolites, it does require an additional time for water quantification which is close to the acquisition time of the water-suppressed data. Ideally, for pure MRS studies, qMRI measurements at this high resolution would be deemed unnecessary, since the spectroscopic voxels are of a much lower resolution. However, accurate water content mapping requires data in high resolution, since the calibration with the CSF voxels is a crucial step and low-resolution images would lead to increased partial volume of CSF voxels and decreased number of CSF voxels in general. Further studies are required to determine the lower limit of the achievable resolution of water content maps without any bias.

## Acknowledgements

We would like to thank Prof. Charles Gasparovic for fruitful discussions on the topic of PSF correction and absolute quantification of metabolites.

