## Supporting Information for "Absolute quantification of cerebral metabolites using 2D ^1^H-MRSI with quantitative MRI-based water reference"

Supporting Informtion Table 1: Literature relaxation times and relative proton densities of water used

|  | WM | GM | CSF |
| --- | --- | --- | --- |
| T1 | 878 ms ^29^ | 1425 ms ^29^ | 4300 ms ^30^ |
| T2 | 58.68 ms ^31^ | 69.45 ms ^31^ | 2000 ms ^32^ |
| H_2_O | 69.99 p.u^29^ | 81.72 p.u^29^ | 100.00 p.u |

Supporting Informtion Table 2: Literature relaxation times of metabolites used ^33^

|  | tNAA | tCr | tCho |
| --- | --- | --- | --- |
| T1 | 1410 ms | 1350 ms | 1190 ms |
| T2 | 271 ms | 154 ms | 197ms |

**PSF correction of qMRI maps and WM, GM and CSF segmentations**

The correction of the difference in the point spread functions between the low-resolution 2D MRSI spectroscopic data and the high-resolution imaging data is described in this section. The correction is specific for Siemens VE11c software version and will be different for data acquired with MRI scanners from other vendors. For the current study, the algorithm for PSF is as follows:

Let the ‘*HR_map’* denote the high-resolution qMRI map/ segmentation. And let ‘*HR_map_resliced’* denote the HR_map resliced into the spectroscopic space.

matrix_size = [X, Y]	# Spectroscopic Matrix

middle_point = [X/2, Y/2]	# Middle point of the matrix

hamm_width = Width of the hamming filter used # default in Siemens = 0.5

data_PSF = zeros(HR_map_resliced.shape)	# Initialize PSF corrected HR image

for i in range(no.of slices (HR_map_resliced*)*):

    data_i = HR_map_resliced [:,:,i]

    data_i(voxels outside VOI) = 0 # Set pixels outside the spectroscopic VOI to 0

```

    inversefft = np.fft.ifftshift(np.fft.ifft2(data_i)) # Inverse fft


    filt = zeros([data_i.shape[0], data_i.shape[1]])

    filt_mod_hamm = zeros(matrix_size)
    ka_image = zeros(matrix_size)

    for x in range(X):
       for y in range(Y):

         k1_max = (x/2 - 0.5)
         k2_max = (y/2 - 0.5)

         delta_k = np.sqrt(((x - middle_point[0])/k1_max)**2 +
                           ((y - middle_point[1])/k2_max)**2)

         ka = (delta_k - (1 - hamm_width))/hamm_width
         ka_image[x,y] = ka

         if ka < 0:
            filt_mod_hamm[x,y] = 1

          else if ka > 1:
             filt_mod_hamm[x,y] = 0.08

          else:
             filt_mod_hamm[x,y] = 0.54 + 0.46*np.cos(np.pi*ka)

          filt[(data_i.shape[0]/2 – X/2): (data_i.shape[0]/2 + X/2),
              (data_i.shape[1]/2 - Y/2): (data_i.shape[1]/2 + Y/2)] = filt_mod_hamm

        inversefft_PSF = inversefft * filt

        data_PSF[:,:,i] = = np.abs(np.fft.fft2(np.fft.fftshift(inversefft_PSF)))
```

**MRSinMRS checklist**

the Minimum Reporting Standards for in vivo- Magnetic Resonance Spectroscopy (MRSinMRS) checklist can be found in Supplementary Table 1.

**Supplementary Table 1**. MRSinMRS checklist for our multi-sequence MRS protocol.

| Goethe University Frankfurt |  |  |  |
| --- | --- | --- | --- |
| 1. Hardware |  |  |  |
| a. Field strength [T] | 3 T | 3 T | 3 T |
| b. Manufacturer | Siemens | Siemens | Siemens |
| c. Model (software version if available) | Prisma (VE11C) | Prisma (VE11C) | Prisma (VE11C) |
| d. RF coils: nuclei (transmit/ receive), number of channels, type, body part | 20 ch ^1^H head coil | 20 ch ^1^H head coil | 20 ch ^1^H head coil |
| e. Additional hardware | N/A | N/A | N/A |
| 2. Acquisition |  |  |  |
| a. Pulse sequence | 2D ^1^H Semi-LASER CSI (vendor-provided) | 2D ^1^H Semi-LASER CSI (vendor-provided, water reference) | ^1^H STEAM SVS (vendor-provided, water reference) |
| b. Volume of Interest (VOI) locations | Healthy volunteer: above the corpus callosum  Patient: tumor and contralateral | Healthy volunteer: above the corpus callosum  Patient: tumor and contralateral | Normal-appearing white matter |
| c. Nominal VOI size [cm^3^, mm^3^] | Adjusted according to tumor volume with a slice thickness of 12 mm | Adjusted according to tumor volume with a slice thickness of 12 mm | 10 x 10 x 10 mm^3^ |
| d. Repetition Time (TR), Echo Time (TE) [ms, s] | TR = 2000 ms, TE = 40 ms | TR = 2000 ms, TE = 40 ms | TR = 10000 ms, TE = 20 ms, TM = 10 ms |
| e. Total number of Excitations or acquisitions per spectrum  In time series for kinetic studies   i. Number of Averaged spectra (NA) per time-point ii. Averaging method (e.g. block-wise or moving average) iii. Total number of spectra (acquired / in time-series) | 2 | 1 | 1 |
| f. Additional sequence parameters  (spectral width in Hz, number of spectral points, frequency offsets)  If STEAM:, Mixing Time (TM)  If MRSI: 2D or 3D, FOV in all directions, matrix size, acceleration factors, sampling method | 2000 Hz, 1024 points  delta frequency = −2.7 ppm  flip angle = 90°  2D: 240 × 240 × 12 mm^3^ FOV; matrix size 20 x 20 interpolated to 32 x 32; no acceleration factor; weighted distribution sampling | 2000 Hz, 1024 points  delta frequency = 0 ppm  flip angle = 90°  2D: 240 × 240 × 12 mm^3^ FOV; matrix size 20 x 20 interpolated to 32 x 32; no acceleration factor; weighted distribution sampling | 1200 Hz, 1024 points  delta frequency = 0 ppm  flip angle = 90° |
| g. Water Suppression Method | CHESS | None | None |
| h. Shimming Method, reference peak, and thresholds for “acceptance of shim” chosen | Automated 3D B0 field mapping technique (GRE-SHIM for brain) | Automated 3D B0 field mapping technique (GRE-SHIM for brain) | Automated 3D B0 field mapping technique (GRE-SHIM for brain) |
| i. Triggering or motion correction method  (respiratory, peripheral, cardiac triggering, incl. device used and delays) | N/A | N/A | N/A |
| 3. Data analysis methods and outputs |  |  |  |
| a. Analysis software | LCmodel 6.2 | LCmodel 6.2 | LCmodel 6.2 |
| b. Processing steps deviating from quoted reference or product | Basis set created using jMRUI 6.0 plug-in NMR-ScopeB | LCModel water reference fit | LCModel water reference fit |
| c. Output measure  (e.g. absolute concentration, institutional units, ratio)Processing steps deviating from quoted reference or product | Ratios to water | Used as water reference for metabolite quantification | Used as water reference for metabolite quantification |
| d. Quantification references and assumptions, fitting model assumptions | The basis set included spectra of N-acetylaspartate, N-acetylaspartylglutamate, glycerophosphocholine, choline, creatine, γ-aminobutyric acid, glucose, glutamate, glutamine, myo-inositol, glutathione, glycine, alanine, lactate, valine. | None | None |
| 4. Data Quality |  |  |  |
| a. Reported variables  (SNR, Linewidth (with reference peaks)) | None | None | None |
| b. Data exclusion criteria | LCModel SNR <3, LCModel FWHM >0.1 ppm. | None | None |
| c. Quality measures of postprocessing Model fitting (e.g. CRLB, goodness of fit, SD of residual) | CRLB < 10% for total choline. | None | None |
| d. Sample Spectrum | None | None | None |
